## Supporting Information, tables and figures for "Seasonally migratory songbirds have different historic population size characteristics than resident relatives"

Table S1. Specimen data and NCBI-SRA numbers (PRJNA1112856). Vouchered specimens are housed in the following institutions: UAM (University of Alaska Museum), MSB (Museum of Southwestern Biology, University of New Mexico), LSUMNS (Louisiana State University Museum of Natural Science), and FMNH (Field Museum of Natural History).

| Institution | Catalog # | Species | Year | Field No. | Locality | NCBI-SRA |
|---|---|---|---|---|---|---|
| UAM | 28620 | *H. mustelina* | 2010 | KSW5403 | Belize: Toledo District; Big Falls | SRR29089747 |
| UAM | 27774 | *C. fuscescens* | 2007 | KSW5151 | Belize: Toledo District; Big Falls | SRR29089742 |
| UAM | 15202 | *C. guttatus* E | 1992 | KSW4013 | USA: Vermont; Brandon | SRR29089748 |
| UAM | 26337 | *C. guttatus* W | 2008 | UAMX5095 | USA: Alaska; Kodiak | SRR29089739 |
| UAM | 22642 | *C. minimus* | 2003 | KSW5000 | USA: Alaska; Fairbanks | SRR29089741 |
| n.a. (blood) | KF15K01 | *C. ustulatus swainsoni* | 2011 | KF15K01 | Canada: British Columbia, Kamloops | pending |
| n.a. (blood) | KF01K01 | *C. u. ustulatus* | 2011 | KF01K01 | Canada: British Columbia, Kamloops | SRS18060177 |
| UAM | 19996 | *C. bicknelli* | 2000 | KSW3633 | USA: Vermont; Mt Mansfield | SRR29089740 |
| UAM | 25341 | *C. aurantiirostris* | 2004 | MJM1154 | Panama: Chiriqui; El Salto | SRR29089749 |
| MSB | 31939 | *C. fuscater* | 2008 | MSB31939 | Peru: Amazonas; 4.5 km N Tullanya | SRR29089746 |
| UAM | 25098 | *C. frantzii* | 2004 | KSW4485 | Panama: Chiriqui; Volcan Baru | SRR29089744 |
| LSUMNS | 138784 | *C. gracilirostris* | 1990 | JMB1065; B-16270 | Costa Rica: San Jose; Cerro de la Muerte | SRR29089750 |
| UAM | 10352 | *C. mexicanus* | 1994 | PEP2489 | Mexico: Veracruz; Volcan San Martin | SRR29089743 |
| FMNH | 343305 | *C. occidentalis* | 1989 | MEX408 | Mexico: Oaxaca; Totontepec | SRR29089745 |

Table S2. Data from PSMC analyses reflecting effective population sizes ($N_e \times 10^4$) through history at depths > 50 Kyr (using variable generation times) and the five variables derived and analyzed from that output. Taxa shaded in gray are Neotropical residents.

| **Taxon** | **Mean $N_e$ (± SD)** | **SD/mean** | **Degree of early growth 1 - ($N_{trough}/N_{peak}$)** | **Rate of early growth degree/deltaT** | **deltaT** |
|---|---|---|---|---|---|
| *Hylocichla mustelina* | 35.63 (± 20.11) | 0.56 | 0.86 | 3.03E-07 | 2,834,197 |
| *Catharus fuscescens* | 95.77 (± 56.07) | 0.58 | 0.83 | 1.79E-07 | 4,627,125 |
| *C. guttatus E* | 78.69 (± 76.09) | 0.97 | 0.90 | 4.89E-07 | 1,850,522 |
| *C. guttatus W* | 40.15 (± 24.11) | 0.60 | 0.81 | 3.59E-07 | 2,266,059 |
| *C. minimus* | 63.71 (± 28.83) | 0.45 | 0.78 | 2.35E-07 | 3,323,013 |
| *C. ustulatus swainsoni* | 76.61 (± 66.61) | 0.87 | 0.79 | 2.12E-07 | 3,740,764 |
| *C. ustulatus ustulatus* | 39.56 (± 12.15) | 0.31 | 0.63 | 2.16E-07 | 2,909,628 |
| *C. bicknelli* | 46.90 (± 24.12) | 0.51 | 0.78 | 3.01E-07 | 2,580,641 |
| *C. aurantiirostris* | 12.39 (± 2.78) | 0.22 | -0.75 | -7.93E-07 | 942,318 |
| *C. fuscater* | 11.33 (± 1.66) | 0.15 | 0.39 | 5.57E-07 | 713,817 |
| *C. frantzii* | 24.97 (± 8.61) | 0.34 | 0.17 | 1.93E-07 | 886,294 |
| *C. gracilirostris* | 30.80 (± 11.08) | 0.36 | 0.58 | 3.25E-07 | 1,775,757 |
| *C. mexicanus* | 37.10 (± 16.18) | 0.44 | 0.75 | 2.92E-07 | 2,575,082 |
| *C. occidentalis* | 76.39 (± 53.14) | 0.70 | 0.91 | 1.97E-07 | 4,595,855 |
| **Means (± SD)** | | | | | |
| migrants | 59.63 (± 38.48)** | 0.61 (± 0.21)* | 0.798 (± 0.08)* | 2.87E-7 (± 1.01E-7) | 3,016,494 (± 875,860)* |
| residents | 32.16 (± 15.58) | 0.36 (± 0.19) | 0.342 (± 0.59) | 1.284E-7 (± 4.71E-7) | 1,914,854 (+ 1,489,246) |

* $p < 0.05$; ** $p < 0.01$

### Supplementary Figures

Figure S1. Distribution maps of the thrush taxa in this study. Among seasonal migrants, green indicates breeding range and yellow is wintering range. Among sedentary lineages (those shaded in gray), purple indicates year-round range. Data are from BirdLife International and the Handbook of the Birds of the World (2021), Bird species distribution maps of the world. Version 2021.1. Available at http://datazone.birdlife.org/species/requestdis.

Figure S2. Graphic presentation of the *Catharus minimus* PSMC dataset, showing effective population size ($N_e \times 10^4$) from 50 kyr back in time to the lineages' origins as estimated from genomic data. The variables in our analyses are the mean and SD of the effective population size ($N_e$) values, *deltaT* of initial growth ($time_{trough} - time_{peak}$), the degree of that growth ($1 - [N_{trough}/N_{peak}]$), and the rate of that growth (degree/*deltaT*).

Figure S3. Montage of the historic effective population size curves of all lineages analyzed in this study, with each lineage in a separate panel (based on pairwise sequentially Markovian coalescent, or PSMC, analyses). Sedentary lineages are highlighted in gray. Note that scales on both axes vary among panels, and that the X axis is on a log scale. Bold red lines are the main curves from the original data, and pink lines reflect 100 replicates from bootstrapped sequences. Bold red curves are all overlaid on common axes in the single panel of Fig. 2.

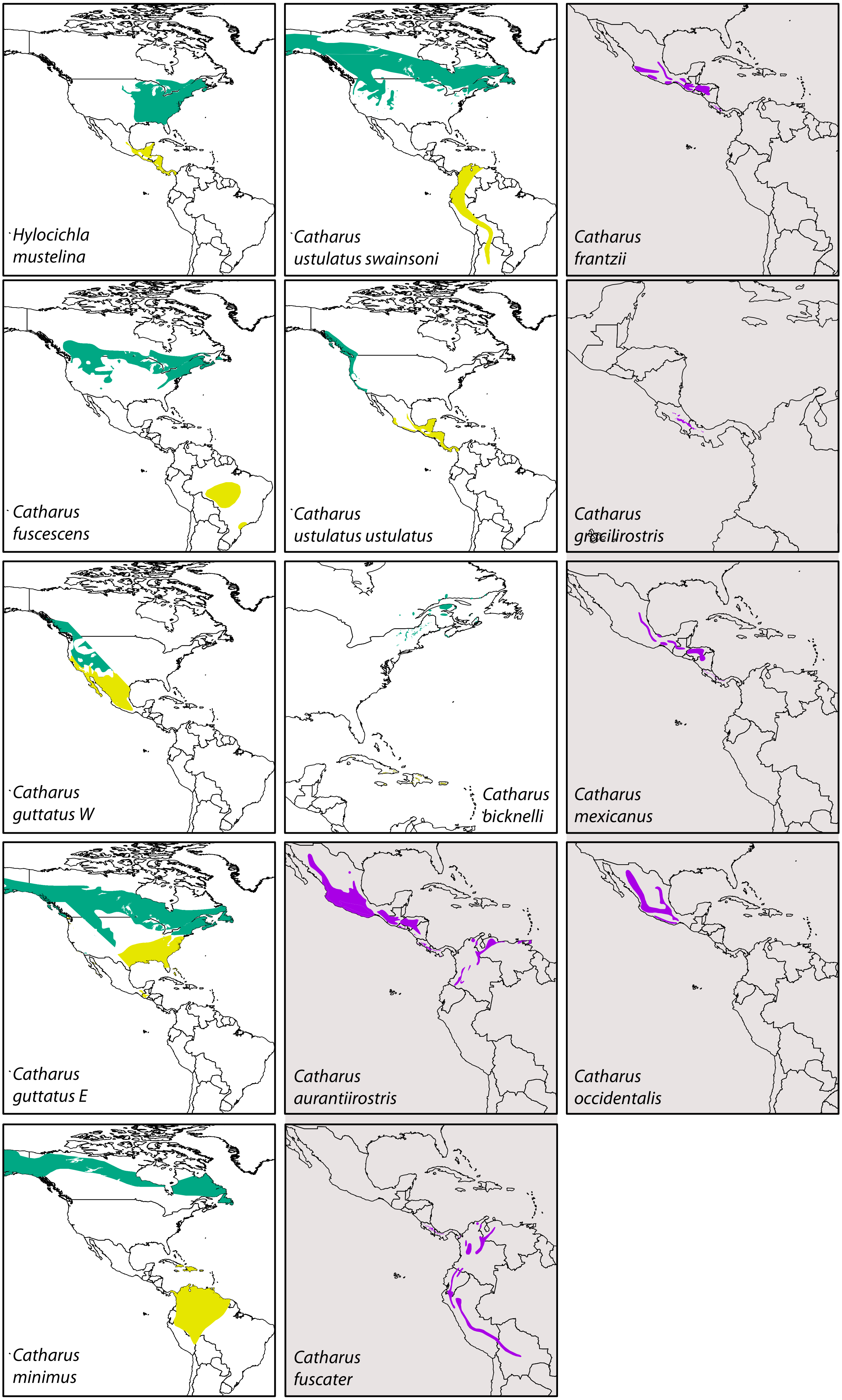

Hylocichla mustelina
Catharus ustulatus swainsoni
Catharus frantzii
Catharus fuscescens
Catharus ustulatus ustulatus
Catharus gracilirostris
Catharus guttatus W
Catharus bicknelli
Catharus mexicanus
Catharus guttatus E
Catharus aurantiirostris
Catharus occidentalis
Catharus minimus
Catharus fuscater

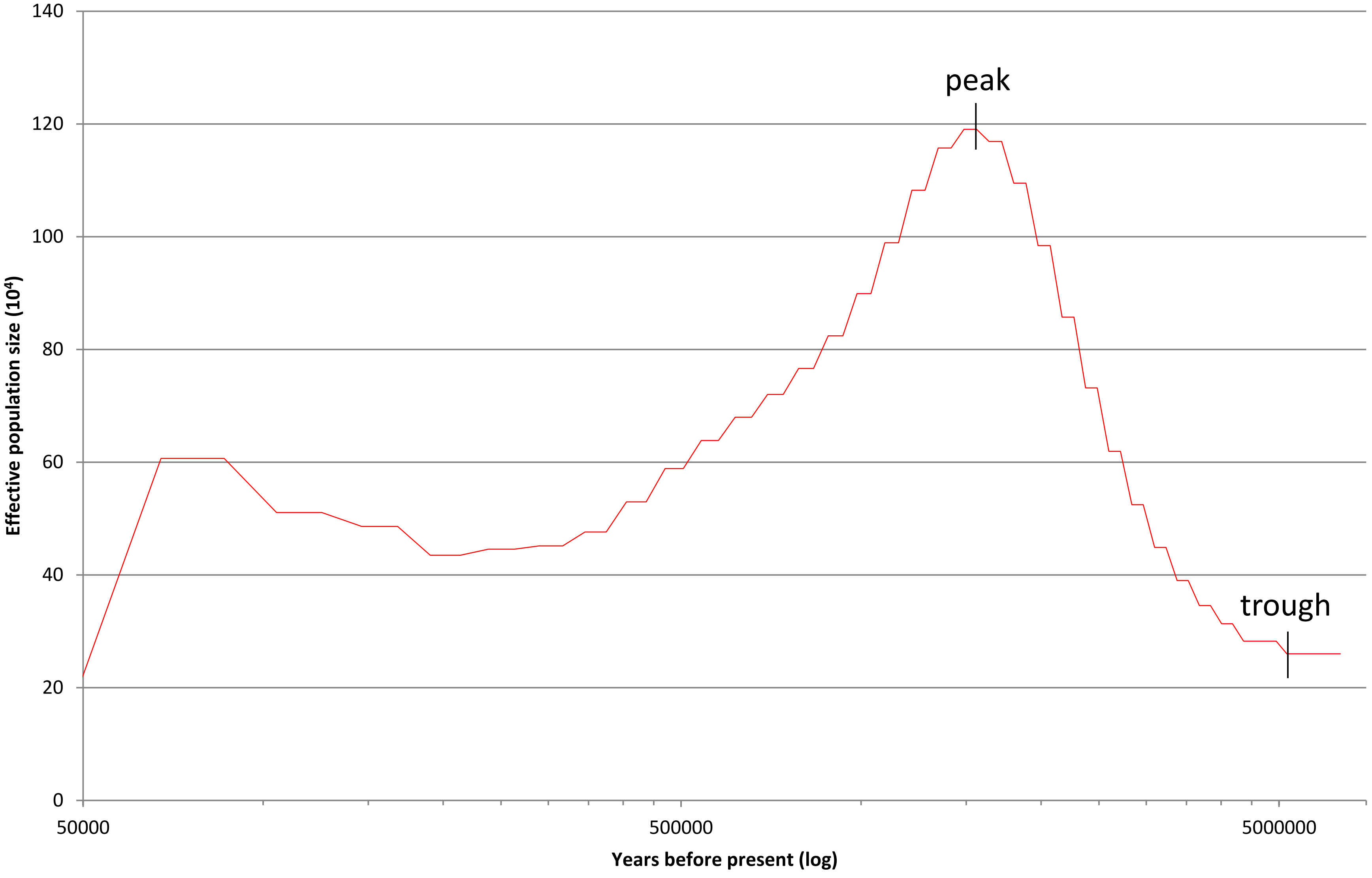


Figure S2. Graphic presentation of the *Catharus minimus* PSMC dataset, showing effective population size ($N_e$) from 50 kyr back in time to the lineages' origins as estimated from genomic data. The variables in our analyses are the mean and SD of the effective population size ($N_e$) values, *deltaT* of initial growth ($time_{trough} - time_{peak}$), the degree of that growth ($1 - [N_{trough}/N_{peak}]$), and the rate of that growth (degree/*deltaT*).

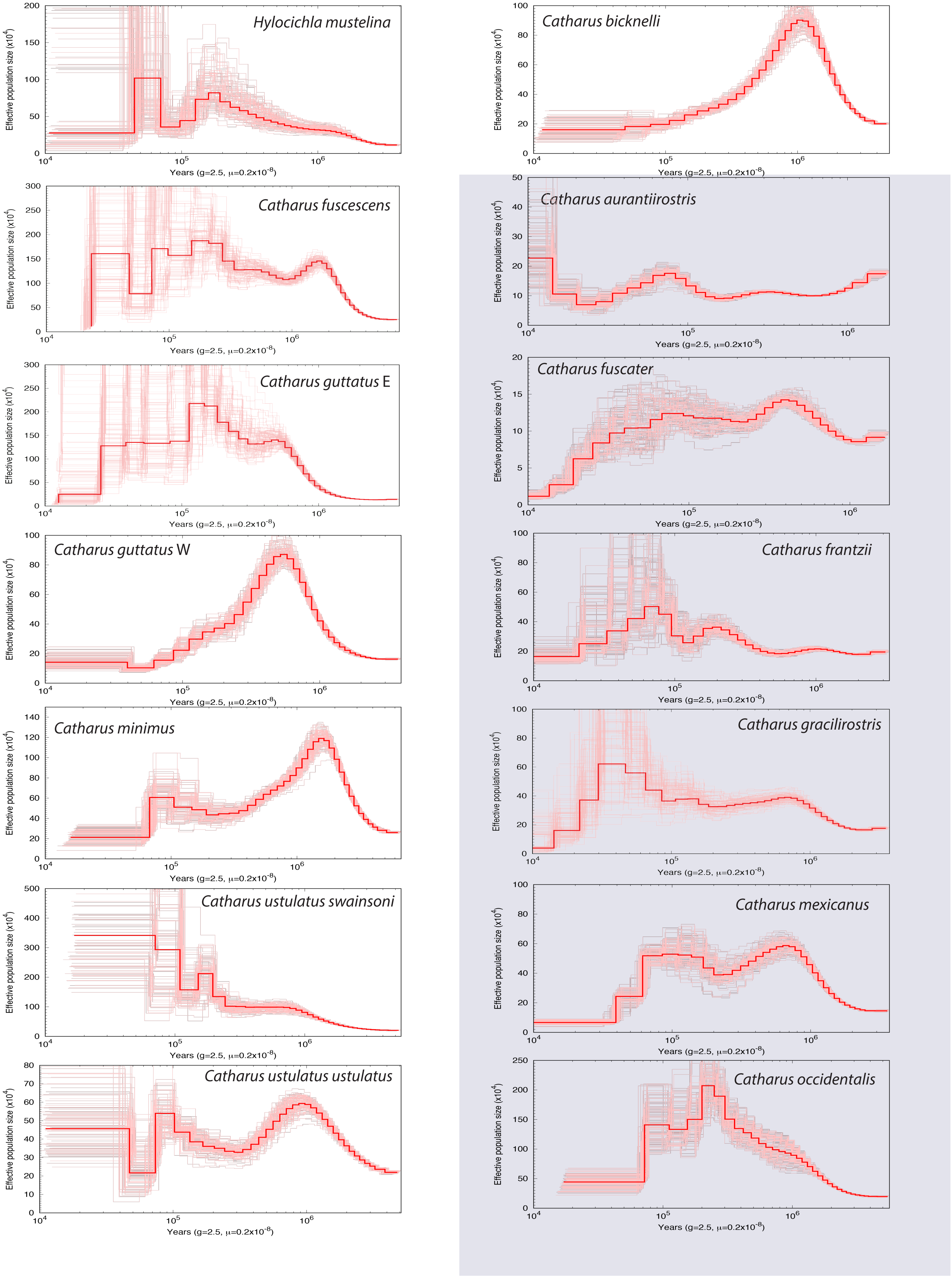
Hylocichla mustelina
Catharus bicknelli
Catharus fuscescens
Catharus aurantiirostris
Catharus guttatus E
Catharus fuscater
Catharus guttatus W
Catharus frantzii
Catharus minimus
Catharus gracilirostris
Catharus ustulatus swainsoni
Catharus mexicanus
Catharus ustulatus ustulatus
Catharus occidentalis
Effective population size (x10^4)
Years (g=2.5, μ=0.2x10^-8)
